# 3.5KJPNv2, An allele frequency panel of 3,552 Japanese Individuals

**DOI:** 10.1101/529529

**Authors:** Shu Tadaka, Fumiki Katsuoka, Masao Ueki, Kaname Kojima, Satoshi Makino, Sakae Saito, Akihito Otsuki, Chinatsu Gocho, Mika Sakurai-Yageta, Inaho Danjoh, Ikuko N. Motoike, Yumi Yamaguchi-Kabata, Matsuyuki Shirota, Seizo Koshiba, Masao Nagasaki, Naoko Minegishi, Atsushi Hozawa, Shinichi Kuriyama, Atsushi Shimizu, Jun Yasuda, Nobuo Fuse, the Tohoku Medical Megabank Project Study Group, Gen Tamiya, Masayuki Yamamoto, Kengo Kinoshita

## Abstract

The first step towards realizing personalized healthcare is to catalog the genetic variations in a population. Since the dissemination of individual-level genomic information is strictly controlled, it will be useful to construct population-level allele frequency panels and to provide them through easy-to-use interfaces.

In the Tohoku Medical Megabank Project, we have sequenced nearly 4,000 individuals from a Japanese population, and constructed an allele frequency panel of 3,552 individuals after removing related samples. The panel is called the 3.5KJPNv2. It was constructed by using a standard pipeline including the 1KGP and gnomAD algorithms to reduce technical biases and to allow comparisons to other populations. Our database is the first largescale panel providing the frequencies of variants present on the X chromosome and on the mitochondria in the Japanese population. All the data are available on our original database at https://jmorp.megabank.tohoku.ac.jp.

## Background & Summary

It is of fundamental importance to catalog the genetic variation in a general population to realize personalized healthcare and personalized medicine. Since different populations show divergent genetic variations, population-specific analyses based on large cohorts are required^1,2^. Since individual-level genomic information is classified as personal data, access to it is controlled strictly. Therefore, allele frequencies have been published in the form of reference panels^3,4^ to clarify population-level differences.

Accordingly, a large allele frequency reference panel based on the genomes of 1,070 Japanese individuals was first published in 2014^5,6^by the Tohoku Medical Megabank (TMM) Project^7^. A subsequent version published which in 2016 included 2,049 individuals^8^, and one distributed in 2017 included 3,554 individuals. These reference panels were used for various other projects. For example, the IRUD project (Japan’s Initiative on Rare and Undiagnosed Diseases^9^) used the reference panel to reduce the discovery of false positive single nucleotide variants (SNVs) during the exome analyses of undiagnosed patients. In another project, CYP SNVs included in the reference panels were selected and analyzed systematically for their effect on drug metabolism^10–12^. As seen from these examples, the previous versions of the reference panels worked well; however, there are some limitations. One of the limitations was the lack of multi-allelic sites as predicted by the infinite site model^13^. Following standard practice, multi-allelic sites were removed from the previous frequency panels, which resulted in a lack of high frequency alleles in the reference panel. Since human genomes are now considered to have accrued a large number of mutations due to a rapid expansion of the population size, the analysis of multi-allelic sites should prove interesting from the view point of human population genetics. However, we hope that it will be described elsewhere. Another limitation with our analysis was the gradual obsolescence of the 1KJPN pipeline. When we constructed 1KJPN, several analysis pipelines were used, but now largescale analyses such as the 1000 genomes project (1KGP)^14^ and the genome aggregation project (gnomAD)^4^ use virtually equivalent protocols for variant calling. The difference in pipelines can make it difficult to compare the allele frequencies of different populations. Thus, we decided to perform a re-analysis of variant calls and construct a new reference panel for the Japanese population. In this paper, we will describe some details of our 1 new panel construction by using a pipeline similar to the 1KGP and gnomAD pipelines. We also report the variant frequencies of X chromosome and those of mitochondria, which makes this the first such report to do so on a largescale for the Japanese population.

## Methods

### Sample Information

Data were obtained from 3,552 individuals in Japan [Table 1]. Among them, 3,342 samples came from individuals who participated in the TMM Project, which was led by the Tohoku Medical Megabank Organization (ToMMo) at Tohoku University and Iwate Tohoku Medical Megabank Organization (IMM) at Iwate Medical University. The TMM project recruited participants from both the Miyagi and Iwate prefectures. Individuals who presumably originated from other prefectures were also included [Table 1(a)]. A further 29 samples came from individuals who participated in the Nagahama Study15. Lastly, 181 samples came from individuals recruited by the National Hospital Organization Nagasaki Medical Center. Written informed consent was obtained from all the participants.

**Table 1.** Summary of samples included in 3.5KJPNv2.

| Participant group | Subtotal |  |
| --- | --- | --- |
| TMM Project |  |  |
| Miyagi or Iwate prefectures* | 1,507 | 3,342 |
| Other than Miyagi and Iwate prefectures | 491 |  |
| Origin information is not available | 894 |  |
| Nagahama Study | 29 |  |
| Individuals recruited in Nagasaki Medical Center | 181 |  |
| **Total** | **3,552** |  |

(b) Participants' age and sex. Age information for 14 participants was missing.
| **Age ranges** | **Male** | **Female** | **Total** |
| --- | --- | --- | --- |
| **29 and younger** | 66 | 173 | 239 |
| **30-39** | 184 | 329 | 513 |
| **40-49** | 124 | 243 | 367 |
| **50-59** | 219 | 402 | 621 |
| **60-69** | 629 | 653 | 1,280 |
| **70-79** | 312 | 194 | 506 |
| **80 and older** | 7 | 5 | 12 |

Participants’ declared age, and their sex as determined from X-and Y-chromosome sequencing are presented in Table 1(b). Samples with irregular karyotypes, such as those with Turner syndrome, were excluded. Close relatives of individual subjects, based on mean identity-by-descent (IBD; PIHAT in PLINK version 1.07) values indicating relatedness closer than between third cousins degree relatives, were excluded.

### Whole-genome sequencing

Library preparation and sequencing were performed as described earlier with minor modifications^5^. Briefly, genomic DNA extracted from the buffy coat was fragmented by sonication to an average target size of 550 bp. After library quantification using the quantitative MiSeq method^15^, sequencing was performed on the HiSeq 2500 system (Illumina). The TruSeq Rapid PE Cluster V1 and SBS Kits (1 sample per flowcell), and the TruSeq Rapid PE Cluster Kit V2 and SBS Kit (2 samples per flowcell) were used for 162-bp and 259-bp paired-end protocols, respectively.

### Whole-genome re-sequencing workflow

We employed a workflow known as the GATK Best Practices workflow, which is becoming the standard procedure globally for whole genome re-sequencing analysis. Several recent large-scale genome analyses, such as the 1000 Genomes project^14^and gnomAD^4^adopted the same workflow. Although we used an original re-sequencing workflow for 1KJPN^5^, 2KJPN, and 3.5KJPNv1, we decided to use a more common pipeline to build 3.5KJPNv2 to allow for comparisons of allele frequencies between different populations. We customized three steps in the GATK Best Practices workflow: (1) the choice of the reference genome, (2) the use of Base Quality Score Recalibration (BQSR), and (3) the joint genotyping step.

### Alignment of sequence reads to the reference genome

FASTQ files of each sample were aligned to a set derived from the human reference genome (GRCh37) that contains the revised Cambridge Reference Sequence (rCRS), un-localized/un-placed contigs, human gammaherpesvirus 4 sequence (NC_007605), and a decoy sequence (hs37d5). Two pseudoautosomal regions (PAR1 and PAR2) on Y chromosome are masked as N. This reference genome sequence is referred to as hs37d5.fa and it is the same reference sequence as that used in the 1000 Genomes project Phase 2 ^14^.

FASTQ files were aligned with hs37d5.fa using BWA-MEM^16^ version 0.7.12 and sorted by their coordinate using the SortSam program included in Picard^17^ version 2.10.6. BWA-MEM was run at “-K 10000000”, in addition to the default options to reduce any differences when we performed calculations with multiple threads. Thereafter, duplicate PCR reads were removed by using the MarkDuplicates command in Picard. The output was written into a BAM (Binary Alignment/Map) format file. Such files will be referred to as the baseline BAM files in this study.

Although the GATK Best Practices workflow recommends that the BQSR step be carried out after the mapping, we did not do so because of the following reasons. Before analyzing our full dataset of 3,552 samples, we evaluated the effect of BQSR on our dataset. For this purpose, we randomly selected 100 samples from our dataset and re-sequenced them using BQSR as described in GATK Best Practices. We also performed resequencing without the BQSR step. Finally, we performed SNP array analyses on both the sets of 100 samples. In other words, we checked concordance among two kinds of genotyping results: (i) genotyping results obtained after the incorporation of BQSR and (ii) results obtained without BQSR. We then performed SNP array analyses, where we used Japonica Array^18^version 1 for the analyses. We did not observe significant differences in concordance between (i) and (ii) according to the markers present in the SNP array (Table 2). In light of the preceding information and given the fact that BQSR requires approximately 10 hours per sample to execute, increasing the total computation time by 50%, we decided to skip the BQSR step in this study.

**Table 2.** Comparison of WGS genotyping and SNP array genotyping.

|  |  | Array genotype |  |  |  |  |
| --- | --- | --- | --- | --- | --- | --- |
|  |  | NotObserved | NoCall | 0/0 | 0/1 | 1/1 |
| **WGS**<br>**(with BQSR)**<br>**genotype** | **NotObserved** | - | 0 | 236 | 4 | 1 |
|  | **NoCall** | 9581 | 0 | 3 | 0 | 0 |
|  | **0/0** | 556282 | 42 | 16301 | 28 | 3 |
|  | **0/1** | 119717 | 33 | 20 | 8905 | 31 |
|  | **1/1** | 95637 | 15 | 2 | 11 | 5339 |

(b) Comparison of WGS genotyping procedure (excluding the BQSR step) and SNP array genotype.
|  |  | Array genotype |  |  |  |  |
| --- | --- | --- | --- | --- | --- | --- |
|  |  | NotObserved | NoCall | 0/0 | 0/1 | 1/1 |
| **WGS**<br>**(without**<br>**BQSR)**<br>**genotype** | **NotObserved** | - | 0 | 236 | 4 | 1 |
|  | **NoCall** | 9791 | 0 | 3 | 0 | 0 |
|  | **0/0** | 552854 | 42 | 16301 | 27 | 3 |
|  | **0/1** | 119012 | 33 | 20 | 8906 | 31 |
|  | **1/1** | 95483 | 15 | 1 | 10 | 5338 |

**Table 3.** Number of variants found on autosomes, X chromosome, and mitochondria.

|  | Autosomes |  | X chromosome |  | X chromosome |  | Mitochondria |
| --- | --- | --- | --- | --- | --- | --- | --- |
|  |  |  | (PAR1, PAR2, XTR) |  | (PAR1, PAR2) |  |  |
|  | Raw | Passed | Raw | Passed | Raw | Passed | Raw |
| SNVs | 51,168,347 | 44,107,909 | 2,065,505 | 1,750,054 | 2,005,093 | 1,726,127 | 2,483 |
| INDELs | 7,283,992 | 5,839,667 | 295,681 | 240,016 | 305,477 | 244,260 | - |
| Multi allelic<br>SNV sites | 1,409,934 | 701,047 | 48,408 | 20,139 | 54,867 | 28,620 | - |

### Variant discovery on autosomes and joint genotyping

Variant calls for each baseline BAM file were made by using the program HaplotypeCaller included in GATK version 3.7, resulting in the generation of Genomic VCF (GVCF) files. These were used to perform multi-sample joint genotyping in the following steps. After generation of GVCFs for all samples, joint genotyping was done using the GenotypeGVCFs program included GATK version 3.7. Joint genotyping of large samples usually takes a large amount of computational time, but it was not feasible for us to perform joint genotyping of 3,500 samples at the same time. To overcome this difficulty, we divided the autosomes into 3 Mb chunks and performed joint genotyping of each chunk across all the samples. After all the chunks were processed, they were concatenated to produce the whole autosome. To avoid edge effects that may be introduced by chunk splitting, we made overlaps of 1 kb between adjacent chunks and checked concordance of variant calls in the overlapping regions. If discordant variant calls were found in the overlapping regions, we removed them. In total, we found 470 discordant variants on the autosomes and they were not included in the results.

After merging of chunks and checking their concordance, we applied the Variant Quality Score Recalibration (VQSR) filter. GATK resource bundle was used as known site information for VQSR step. We opted for SNV filtration based on the following VQSR scores: QD (variant call confidence normalized by depth of sample reads supporting a variant), MQ (root-mean-square value of the mapping quality of reads across all samples), MQRankSum (rank-sum test for mapping qualities of REF versus ALT reads), ReadPosRankSum (rank-sum Test for relative positioning of REF versus ALT alleles within reads), FS (strand bias estimated using Fisher’s exact test), SOR (strand bias estimated using the symmetric odds ratio test), DP (total depth of coverage per sample and over all samples), and InbreedingCoeff (likelihood-based test for the inbreeding among samples). For INDEL filtration, we excluded the MQ and MQRankSum scores from the preceding list. Finally, we collected the SNVs and INDELs that passed the VQSR step. The numbers of SNVs and INDELs found on the autosomes and the X chromosome are shown in Table. 3.

### Variant discovery on X chromosome

The difference between the analyses of the autosomes and those of the sex chromosomes was the ploidy settings for calling single-sample variants during the GVCF file generation stage. It is well known that there are pseudoautosomal regions (PAR) on the X and Y chromosomes that have similar sequences. Thus, variant calling for these regions should be done with different ploidy settings according to the sex of each sample. The sex of each sample was determined using PLINIK (1.9b05). Since the female samples have two X chromosomes, we treated their reads as having originated from the diploid genome and performed variant calling for them just as we did for the autosomes. For the male samples that have one X chromosome and one Y chromosome, we performed variant calling for PAR and non-PAR reads using different ploidy settings. We treated the PAR reads as having originated from the diploid genome, and non-PAR reads as having originated form the haploid genome. The existence of at least two PARs, called PAR1 and PAR2, has been accepted by most international genome projects. However, there is some discussion about the existence of another similar pseudoautosomal region called the X-transposed region (XTR)^19–21^. To check the pseudoautosomal nature of XTR, we observed heterozygosity of X chromosome for male samples by SNP array analyses. We found significantly higher heterozygosity in three regions, including the XTR in the Japanese population (Fig. 1). Therefore, we decided to use two different SNV calling procedures: one where only PAR1 and PAR2 were considered and the other where XTR was also treated as a pseudoautosomal region. After the genotyping step, we extracted the unmapped reads and the reads that were mapped onto the X and Y chromosomes from the baseline BAM files and then re-mapped them onto a modified version of hs37d5.fa in which XTR along with two pseudoautosomal regions on Y chromosome were masked as N. Except for the mapping step, we employed the same variant calling workflow as we did for the autosomes and X chromosome and considered both PAR1 and PAR2.

**Fig. 1:**
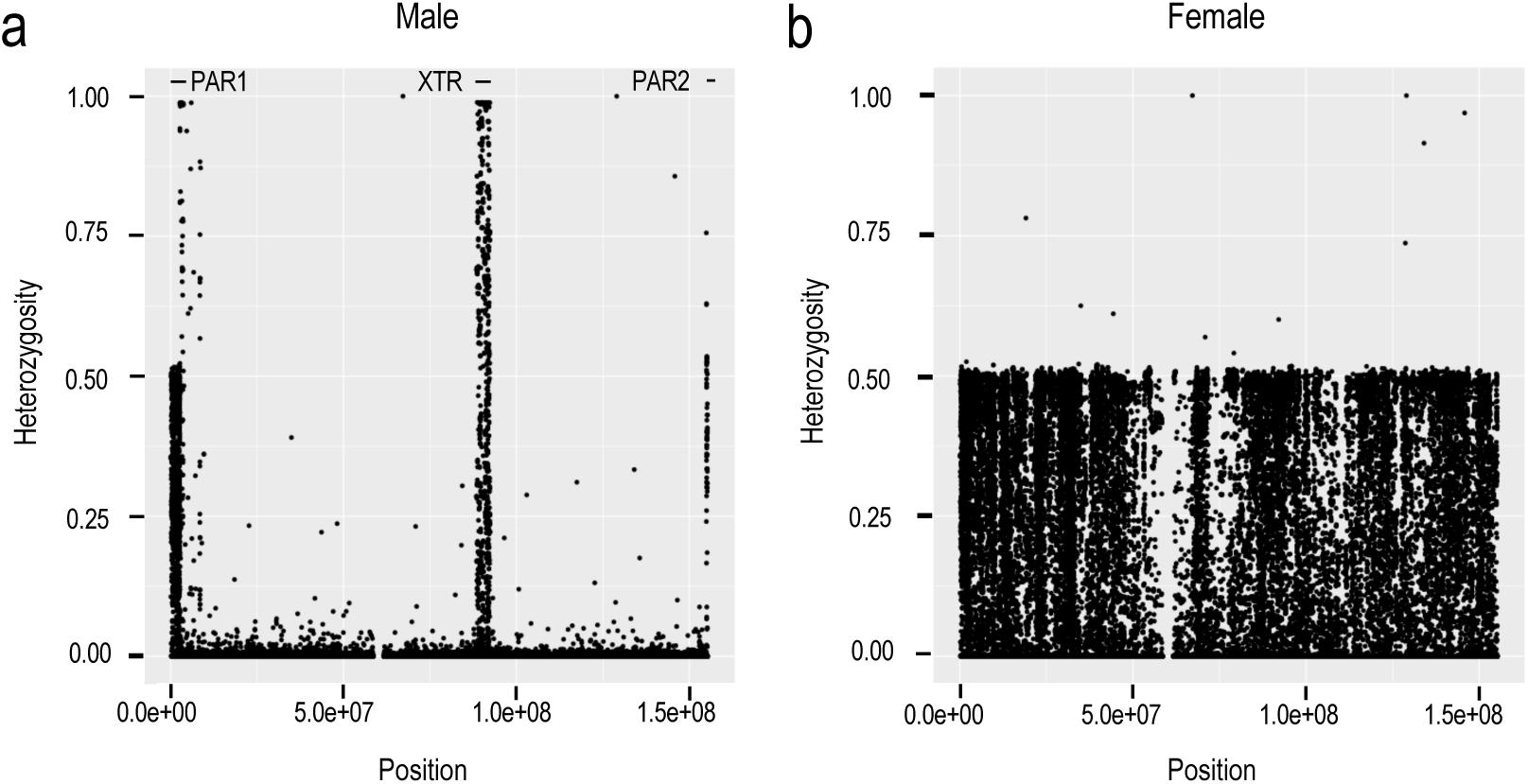
Heterozygosity of X chromosome observed by SNP array analysis.

We constructed X chromosome allele frequency panels for the two ploidy settings by generating the GVCF files and by joint genotyping as described before. We also used GATK resource bundle as known site information for VQSR step.

### Variant discovery on mitochondrial DNA

Since the mitochondrial DNA is circular, we analyzed data originating from the mitochondrial genomes by converting the circular DNA sequence into two linear DNA sequences by inserting a breakpoint within it. One of the linear DNA sequences was the same as used by rCRS, while the other one was generated by shifting the breakpoint by 10,000 bases. The shift was introduced to avoid any edge effects at the breakpoint on the variant calls. First, we extracted unmapped reads and reads aligning onto mitochondrial genome from the baseline BAM files. Thereafter, we re-aligned them onto the two linear mitochondrial genomes using BWA MEM version 0.7.12. Afterwards, we used GATK HaplotypeCaller version 3.7 to detect the variants. Mitochondrial genomes are known to be heteroplasmic. However, we ignored this consideration while building the current version of the variant panel because our focus was on determining the major variants in the first step of analyses for the Japanese population. Therefore, we treated the mitochondrial DNA as haploid when we performing the variant calls.

### Variant annotation

Variants found from 3,552 individuals were annotated by snpEff^22^ version 4.3t while using GENCODE^23^ release 28. GENCODE release 28 is not provided as a pre-built database for snpEff. Therefore, we manually converted the GTF file downloaded from the GENCODE website to a snpEff database according to the instructions given in the snpEff online manual. In addition to GENCODE gene annotations, rs numbers of each variant were resolved by SnpSift^24^ version 4.3t while using dbSNP^25^release 150.

### Code availability

Code written by us especially for this analysis is available on GitHub: https://github.com/gpc-gr/panel3552-scripts. Third-party software employed in this workflow is described in the Methods section.

## Data Records

3.5KJPNv2 is available at the Japanese Multi Omics Reference Panel (jMorp)^26^with web interface (https://jmorp.megabank.tohoku.ac.jp/201901/), and the raw data in VCF (Variant Call Format) format was registered at the NBDC Human Database (https://humandbs.biosciencedbc.jp/en/) with accession code hum0015.v3 by the National Bioscience Database Center (NBDC) of the Japan Science and Technology Agency (JST) to ensure accessibility, preservation and stability of the 3.5KJPNv2 datasets.

Individual’s sequence data and genotyping results from which allele frequency dataset is constructed and validated are available upon request after approval of the Ethical Committee and the Materials and Information Distribution Review Committee of Tohoku Medical Megabank Organization.

## Technical Validation

### Genotype concordance between WGS and SNP array

Out of all the samples, 3,399 samples included in 3.5KJPNv2 had both whole genome sequence data and results obtained after genotyping according to the SNP array available for them. We confirmed the concordance between genotypes obtained by re-sequencing and genotypes obtained by the SNP array (Fig. 2), the result showed that most samples have a high concordance (>= 99.0%).

**Fig. 2:**
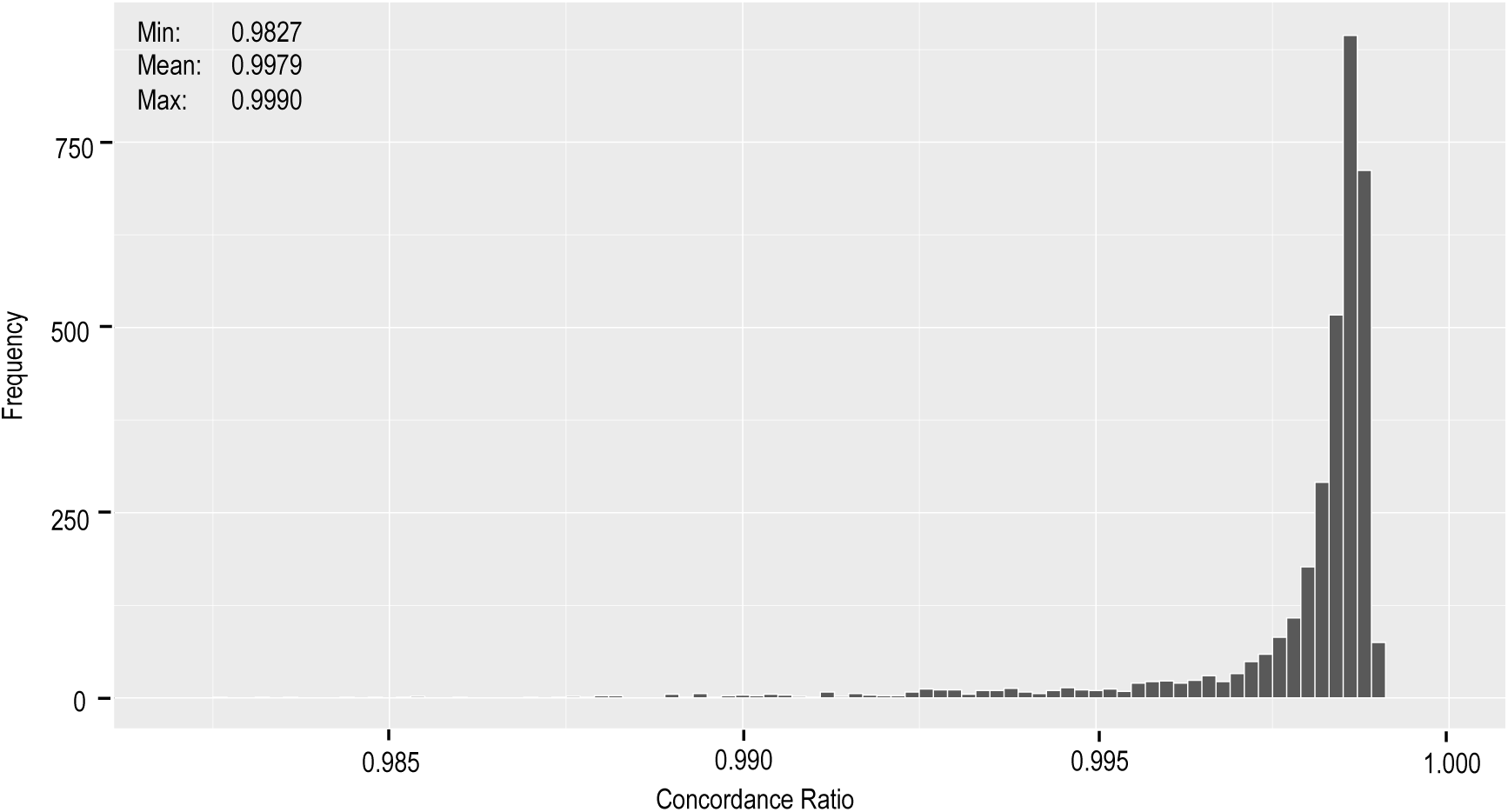
Concordance between re-sequencing result and SNP array genotype.

### Comparison with other Genomes data

We compared the allele frequencies obtained by us as part of 3.5KJPNv2 to those of the whole East Asian population (EAS) obtained by the genome aggregation database gnomAD project. Here, we have shown the results for chromosome 6. As seen in Fig. 3(a), the allele frequencies of SNVs in the two populations correlate highly as expected (Pearson correlation coefficient = 0.829). On the other hand, we could also observe some outliners. Table 4(a) describes several outliers falling in the regions marked as (i), (ii) and (iii) in Fig. 3(a). Region (i) contained five outliners, two of which were located in low complexity regions as identified by RepeatMasker (4.0.0; http://www.repeatmasker.org/). A previous study^27^ suggested that the complexity of the genome sequence can affect the accuracy of short-read aligners, and thus it would be difficult to perform short-read sequencing analyses. Region (ii) and (iii) contained approximately 20 SNVs in total and all of which were located around the HLA region (HLA-DQA1 gene and HLA-DQB1 gene). Again, these are known to be difficult regions for short-read analyses due to their high diversity^28,29^. In both cases, we think that most of the outliers resulted because of poor alignment of the short reads. Some of these outliers may be resolved upon re-analysis with next-generation long-read sequencers.

**Table 4.** Overview of outliers found in allele frequency comparison plots.

|  | Position | Ref/Alt | 3.5KJPNv2 | gnomAD EAS | Possible reason |
| --- | --- | --- | --- | --- | --- |
| Fig. 2(a) – (i) | 3259463 | G/T | 0.6337 | 1.0000 | Unknown |
|  | 25452223 | T/C | 0.3705 | 0.8483 | Low complexity region |
|  | 35283958 | T/C | 0.6488 | 1.0000 | Low complexity region |
|  | 88051052 | T/C | 0.5943 | 1.0000 | Unknown |
|  | 166721424 | C/T | 0.4401 | 0.8750 | Unknown |
| Fig. 2(a) – (ii) | 32609253 | G/A | 0.5351 | 0.0777 | HLA region (HLA-DQA1) |
|  | (15 SNVs omitted) |  |  |  |  |
|  | 32629146 | G/A | 0.5370 | 0.0037 |  |
| Fig. 2(a) – (iii) | 32609379 | C/T | 0.7194 | 0.2405 | HLA region (HLA-DQB1) |
|  | 32610825 | A/G | 0.7245 | 0.2357 |  |
|  | 32629257 | T/A | 0.7793 | 0.2308 |  |
|  | 32629161 | A/G | 0.7201 | 0.0683 |  |
|  | 32629193 | C/T | 0.7211 | 0.0293 |  |
|  | 32629247 | A/C | 0.7171 | 0.0157 |  |

(b) Summary of outliers found in Fig. 3(b).
|  | Position | Ref/Alt | 3.5KJPNv2 | gnomAD EAS | Possible reason |
| --- | --- | --- | --- | --- | --- |
| **Fig. 2(b) – (iv)** | 93743452 | A/T | 0.3535 | 0.1842 | Unknown<br>(gnomAD EAS = 0.3532) |

**Fig. 3:**
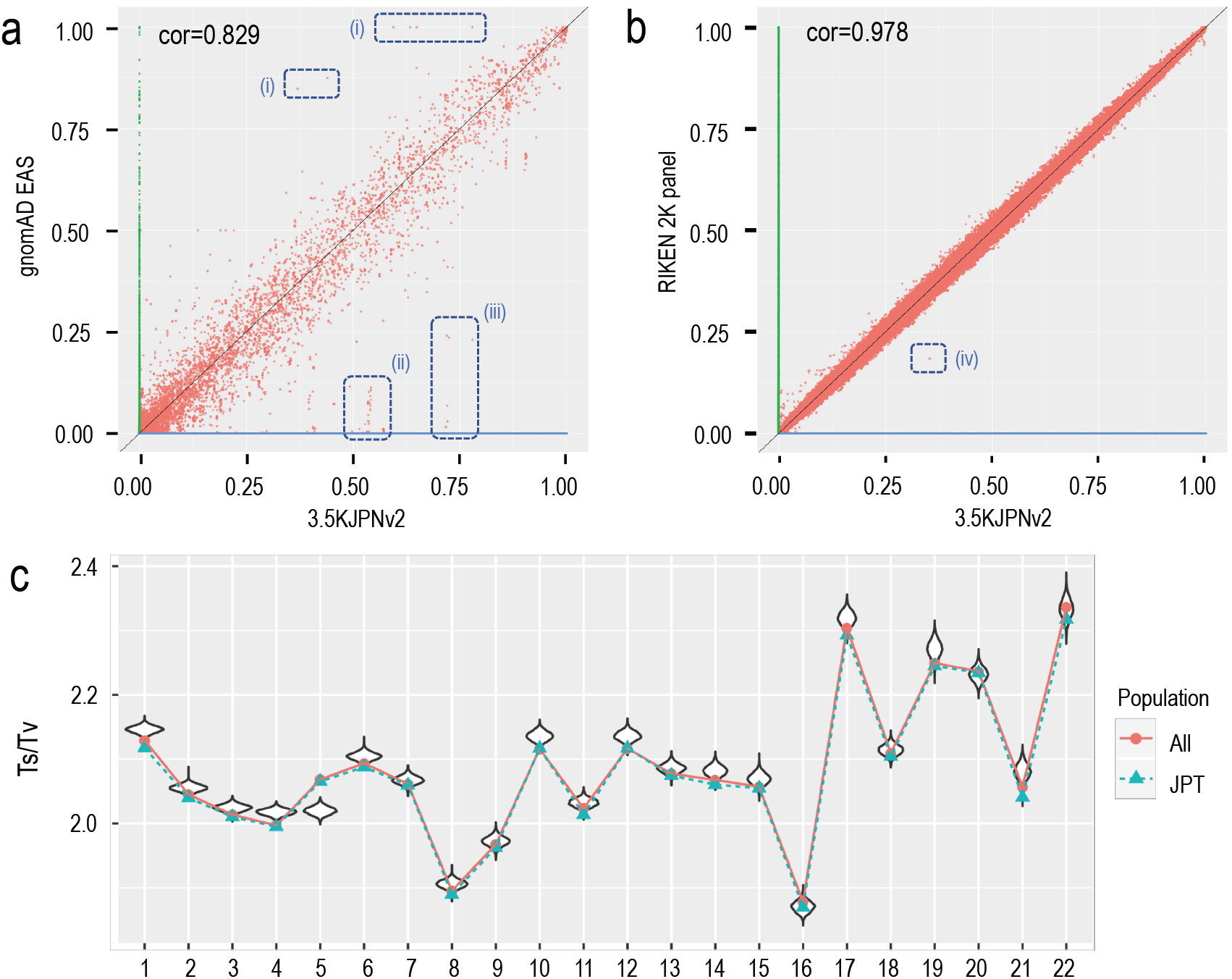
Comparisons of 3.5KJPNv2 with other Genomes data.

We also compared our reference panel with the RIKEN reference panel^30^ consisting of 2000 Japanese individuals, independent of our samples [Fig. 3(b)]. On comparing Fig. 3(a) and Fig. 3(b), we can see a higher consistency between 3.5KJPNv2 and the RIKEN reference panel, although some outliners also exist in Fig. 3(b). There are two main differences between our panel and the RIKEN panel. The first difference is at the filtration step for generating variant calls. We used VQSR for variant filtering after genotyping, while the RIKEN panel used several hard filters in addition to VQSR. We don’t insist that VQSR is better than some combination of hard filtering, but we employed VQSR to reduce any bias introduced by pipeline differences. Another difference is that our panel was constructed for a general population, while RIKEN panel was generated based on patient volunteers. We are not sure that this difference would have a large impact on allele frequencies of most SNVs. However, we think it would be important to consider this difference when our panel is used for personalized healthcare. In other words, allele frequencies of some rare variants can change due to this difference, which in turn can cause large differences in the ability to identify disease-causing variants and to evaluate genetic risks.

We also checked distribution of Ts/Tv metrics for 3.5KJPNv2 and for the 1000 Genomes for each chromosome. In Fig. 3(c) the horizontal axis represents the chromosomes and the vertical axis shows the ratio of transitions and transversions (Ts/Tv ratio). Violin plots indicate the distributions of Ts/Tv ratios among individuals included in 3.5KJPNv2. The red dots represent Ts/Tv ratios for each chromosome compared against all samples the East Asian genomes in the 1000 Genomes project, while the green dots represent comparison against the JPT samples (Japanese samples taken from Tokyo, Japan) included in the 1000 Genomes project. As a result, most of the Ts/Tv values, except for the ones obtained for chromosome 5, were highly like those obtained by the 1000 Genomes project, though slightly higher.

### Analysis of population structure of 3.5KJPNv2

To observe the population structure in 3.5KJPNv2, we created a PCA plot for individuals included in 3.5KJPNv2 and the East Asian populations included in the 1000 Genomes Project. For the East Asian populations, we used the 1000 Genomes Project Phase 3 genotype data, available in the VCF format, for the following populations: CHB (Han Chinese in Beijing, China), JPT (Japanese in Tokyo, Japan), CHS (Southern Han Chinese), CDX (Chinese Dai in Xishuangbanna, China), and KHV (Kinh in Ho Chi Minh City, Vietnam). We obtained a combined genotype dataset by converting the genotype dataset of 3.5KJPNv2 and the dataset of East Asian populations by PLINK. During the conversion, variants with MAF < 0.01 or HWE < 1.0e-5 were removed. The two resultant BED files were combined on commonly existing variants. For the combined dataset, we removed variants with MAF < 0.05, HWE <0.05, or those with a missing rate > 0.01. PCA was applied after LD pruning for the remaining variants with PLINK by selecting the “--indep-pairwise 200 4 0.1” option. We used the same PLINK parameters as Nagasaki et al., 2015 ^5^.

In the PCA plot [Fig. 4(a)], the East Asian populations CHB, CHS, KHV, and CDX populations got aligned in a line reflecting their geographical relationship. The 3.5KJPNv2 individuals and the JPT population overlapped with each other and formed a separate cluster from the CHB, CHS, KHV, and CDX populations. Although there existed another small separate cluster of 12 individuals in the bottom left part of the larger cluster of 3.5KJPNv2 individuals, we could not observe high pairwise IBD values among these individuals. These were at most 0.0223 IBD according to estimation using the PLINK “--genome” option. In addition, these 12 individuals did not form a cluster in the PCA plot of only 3.5KJPNv2 individuals [Fig. 4(b)].

**Fig. 4:**
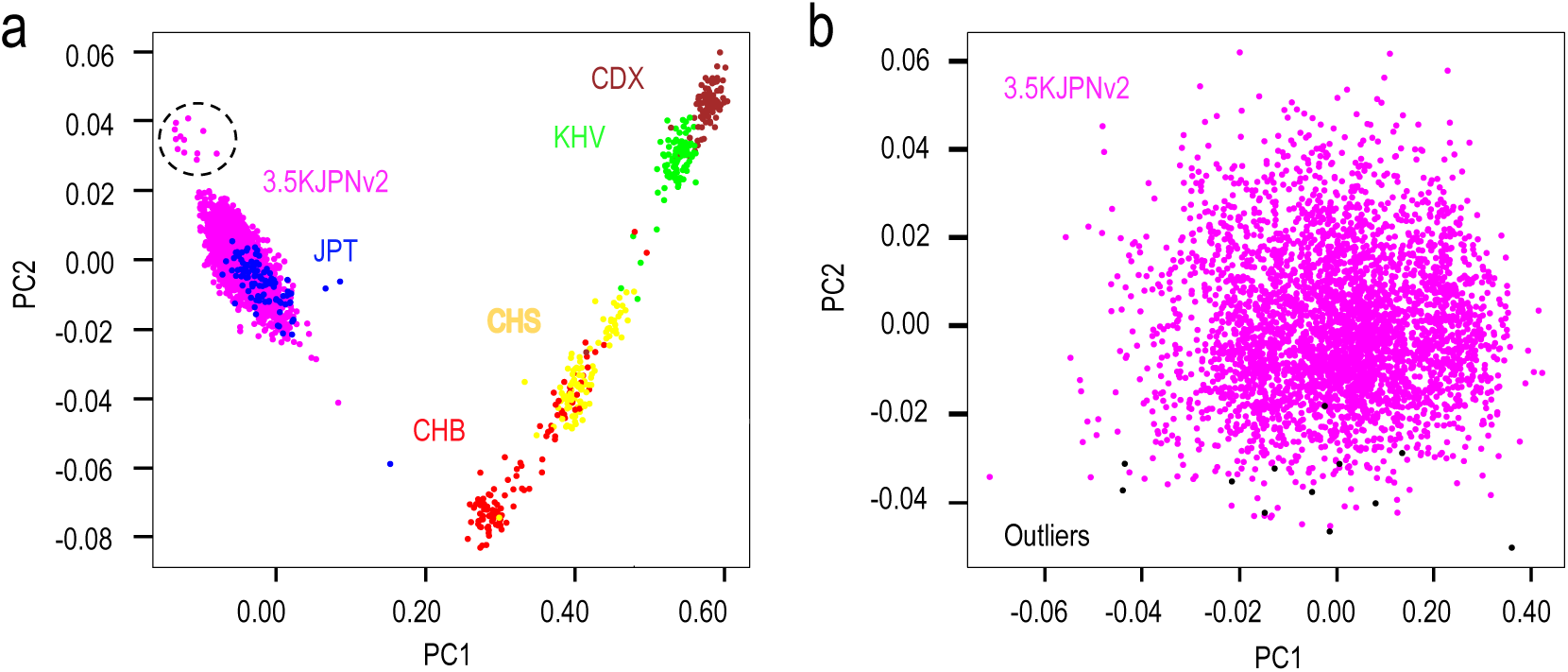
Population structure of 3.5KJPNv2.

## Usage Notes

### Availability of 3.5KJPNv2 allele frequency panel with web interface

3.5KJPNv2 is distributed from jMorp (Japanese Multi Omics Reference Panel) with web interface. jMorp was originally published as a database of metabolites and proteins in plasma obtained from volunteers in ToMMo, which was already described by Tadaka et al^26^. From jMorp release 201806 (Jun 2018, https://jmorp.megabank.tohoku.ac.jp/201806/), genomic variant data has been added, and the latest version 201901 (Jan 2019, https://jmorp.megabank.tohoku.ac.jp/201901/) provides, where allele frequencies of all the genomic variants can be examined through the web interface. Adding genomic variant information enhances multi-layer omics analysis further. Details of the web usage is described in the tutorial section of the web page at https://jmorp.megabank.tohoku.ac.jp/201901/help/tutorial.

## Acknowlegements

We appreciate all the volunteers who participated in the ToMMo project. This work was supported in part by the Tohoku Medical Megabank Project from the Ministry of Education, Culture, Sports, Science and Technology (MEXT) and the Reconstruction Agency; the Japan Agency for Medical Research and Development (AMED; Grant Numbers JP18km0105001 and JP18km0105002) for Tohoku University and (Grant Numbers 18km0105003 and 18km0105004) for Iwate Medical University. All computational resources were provided by the Tohoku University Tohoku Medical Megabank Organization (ToMMo) supercomputer system (http://sc.megabank.tohoku.ac.jp/en), which is supported by Facilitation of &D Platform for AMED Genome Medicine Support conducted by AMED (Grant Number JP18km0405001). We thank Mr. Kota Jin and Mr. Shun Kodate for the web design.

## Author contributions

FK, MN, NM, AH, SK, AS, JY, NF, GT, MY, and KKi designed the study. FK, MN, NM, AH, SK, JY, and NF contributed to sample collection. FK, SS, and AO contributed to sequence data collection. MSY, ID, and GT contributed to array data collection. ST, MU, SM, KKo, GT, and KKi performed computational analysis. INM, YYK, MS, SK, AS, JY, NF, GT, MY, and KKi contributed to data interpretation. ST, INM, YYK, MS, SK, and KKi constructed the jMorp website. ST, FK, KKo, and KKi wrote the initial draft of the manuscript. All authors discussed the results, and approved the final version of the manuscript.

## Competing interests

No competing interests declared.

## References

1. Sudlow, C. et al. UK Biobank: An Open Access Resource for Identifying the Causes of a Wide Range of Complex Diseases of Middle and Old Age. PLoS Med. 12, (2015).

2. Scholtens, S. et al. Cohort Profile: LifeLines, a three-generation cohort study and biobank. Int. J. Epidemiol. 44, 1172–80 (2015).

3. Tennessen, J. A. et al. Evolution and functional impact of rare coding variation from deep sequencing of human exomes. Science (80-.). 336, 64–69 (2012).

4. Lek, M. et al. Analysis of protein-coding genetic variation in 60,706 humans. Nature 536, 285–291 (2016).

5. Nagasaki, M. et al. Rare variant discovery by deep whole-genome sequencing of 1,070 Japanese individuals. Nat. Commun. 6, 1–13 (2015).

6. Yamaguchi-Kabata, Y. et al. iJGVD: an integrative Japanese genome variation database based on whole-genome sequencing. Hum. Genome Var. 2, 15050 (2015).

7. Kuriyama, S. et al. The Tohoku Medical Megabank Project: Design and Mission. J. Epidemiol. 26, 493–511 (2016).

8. Yamaguchi-Kabata, Y. et al. Evaluation of reported pathogenic variants and their frequencies in a Japanese population based on a whole-genome reference panel of 2049 individuals article. J. Hum. Genet. 63, 213–230 (2018).

9. Adachi, T. et al. Japan’s initiative on rare and undiagnosed diseases (IRUD): Towards an end to the diagnostic odyssey. Eur. J. Hum. Genet. 25, 1025–1028 (2017).

10. Hishinuma, E. et al. Functional characterization of 21 allelic variants of dihydropyrimidine dehydrogenase identified in 1070 Japanese individuals. Drug Metab. Dispos. 46, 1083–1090 (2018).

11. Watanabe, T. et al. Functional characterization of 40 CYP2B6 allelic variants by assessing efavirenz 8-hydroxylation. Biochem. Pharmacol. 156, 420–430 (2018).

12. Kumondai, M. et al. Development and application of a rapid and sensitive genotyping method for pharmacogene variants using the single-stranded tag hybridization chromatographic printed-array strip (STH-PAS). Drug Metab. Pharmacokinet. (2018). doi:10.1016/j.dmpk.2018.08.003

13. Kocher, T. D. & Wilson, A. C. Sequence evolution of mitochondrial DNA in humans and chimpanzees: control region and a protein-coding region. in Evolution of Life Fossils, Molecules, and Culture 45, 391–413 (Springer Japan, 1991).

14. Auton, A. et al. A global reference for human genetic variation. Nature 526, 68–74 (2015).

15. Katsuoka, F. et al. An efficient quantitation method of next-generation sequencing libraries by using MiSeq sequencer. Anal. Biochem. 466, 27–29 (2014).

16. Li, H. Aligning sequence reads, clone sequences and assembly contigs with BWA-MEM. 00, 1–3 (2013).

17. Broad Institute. Picard tools. https://broadinstitute.github.io/picard/ (2016).

18. Kawai, Y. et al. Japonica array: Improved genotype imputation by designing a population-specific SNP array with 1070 Japanese individuals. J. Hum. Genet. 60, 581–587 (2015).

19. Cotter, D. J., Brotman, S. M. & Wilson Sayres, M. A. Genetic diversity on the human X chromosome does not support a strict pseudoautosomal boundary. Genetics 203, 485–492 (2016).

20. Skaletsky, H. et al. The male-specific region of the human Y chromosome is a mosaic of discrete sequence classes. Nature 423, 825–837 (2003).

21. Ross, M. T. et al. The DNA sequence of the human X chromosome. Nature 434, 325–337 (2005).

22. Cingolani, P. et al. A program for annotating and predicting the effects of single nucleotide polymorphisms, SnpEff: SNPs in the genome of Drosophila melanogaster strain w1118; iso-2; iso-3. Fly (Austin). 6, 80–92 (2012).

23. Harrow, J. et al. GENCODE: The reference human genome annotation for the ENCODE project. Genome Res. 22, 1760–1774 (2012).

24. Cingolani, P. et al. Using Drosophila melanogaster as a model for genotoxic chemical mutational studies with a new program, SnpSift. Front. Genet. 3, (2012).

25. Sherry, S. T. dbSNP: the NCBI database of genetic variation. Nucleic Acids Res. 29, 308–311 (2001).

26. Tadaka, S. et al. JMorp: Japanese Multi Omics Reference Panel. Nucleic Acids Res. 46, D551–D557 (2018).

27. Phan, V., Gao, S., Tran, Q. & Vo, N. S. How genome complexity can explain the difficulty of aligning reads to genomes. BMC Bioinformatics 16, (2015).

28. Szolek, A. et al. OptiType: Precision HLA typing from next-generation sequencing data. Bioinformatics 30, 3310–3316 (2014).

29. Nariai, N. et al. HLA-VBSeq: Accurate HLA typing at full resolution from whole-genome sequencing data. BMC Genomics 16, (2015).

30. Okada, Y. et al. Deep whole-genome sequencing reveals recent selection signatures linked to evolution and disease risk of Japanese. Nat. Commun. 9, (2018).

